# Preadipocyte uncoupling protein 1 expression invoked by fibroblast growth factors imprints on post-differentiation white and brown adipocyte function

**DOI:** 10.1101/2025.06.26.661774

**Authors:** Katharina Kuellmer, Carolin Mönch, Wilma Roch, Christine Wurmser, Brice Emanuelli, Patricia S. S. Petersen, Hildigunnur Hermannsdóttir, Mersiha Hasic, Tobias Fromme

**Author notes:** Correspondence: Tobias Fromme, Chair of Molecular Nutritional Medicine, TUM School of Life Sciences, Technical University of Munich, Gregor-Mendel-Str. 2, 85354-Freising, Germany.

## Abstract

The presence of uncoupling protein 1 (Ucp1) is a hallmark of thermogenic adipocytes and enables heat production by dissipating energy from mitochondrial proton motive force as heat. The purpose of its recently discovered presence in preadipocytes in response to certain fibroblast growth factors remains elusive. In this study, we systematically investigated the potential of all paracrine fibroblast growth factors (FGFs) to invoke *Ucp1* expression in murine preadipocytes derived from interscapular brown and inguinal white adipose tissue. The factors FGF2, FGF4, FGF8, and FGF9 induced Ucp1 expression in undifferentiated preadipocytes, with FGF2 acting most potently and rapidly. This premature Ucp1 induction did not translate into increased Ucp1 protein abundance or thermogenic activity after full adipogenic differentiation. Notably, preadipocyte treatment with FGFs and concomitant Ucp1 expression led to a sustained suppression of interferon-stimulated genes after differentiation. Preadipocyte Ucp1 was required and sufficient for this lasting imprint. Thus, Ucp1 in preadipocytes programs post-differentiation inflammatory status.

## Introduction

Uncoupling protein 1 (Ucp1) is a protein of the mitochondrial inner membrane and central mechanistic component of non-shivering thermogenesis (Matthias et al., 2000). Activated by free fatty acids, it uncouples oxygen consumption from ATP synthesis by mediating a proton flux from the intermembrane space into the mitochondrial matrix. The expression of Ucp1 is thought to be strictly limited to thermogenic brown or beige adipocytes (Ricquier, 2011). The former are found within the specialized mammalian heater organ brown adipose tissue (BAT), the latter interspersed within otherwise white adipose tissue depots (Wu et al., 2012). The recruitment and differentiation of these cell types is of considerable importance. The efficacy of potential drugs targeting non-shivering thermogenesis as a means to combat metabolic disease in humans would be limited by the small total amount of brown and beige fat, typically in the range of 300g in an adult healthy human (Gerngross et al., 2017; Virtanen et al., 2009).

We and others previously reported the surprising discovery of Ucp1 in non-differentiated preadipocytes in response to treatment with fibroblast growth factors (FGF) 6, 8 and 9 (Gantert et al., 2021; Shamsi et al., 2020; Westphal et al., 2019). These FGFs are members of an ancestral subgroup of FGFs featuring a heparin anchor, rendering their mode of action to be paracrine signaling as opposed to the widely known endocrine FGFs, e.g. FGF21 (Ornitz et al., 1996; Ornitz & Itoh, 2015, 2022). Paracrine FGFs bind FGF receptors (FGFRs) with heparin as a cofactor instead of the obligatory Klotho co-receptors of endocrine FGFs (Kurosu et al., 2006; Urakawa et al., 2006; Yayon et al., 1991). They signal through one or several of four FGF receptors with FGF-specific affinity patterns (FGFR1-4), with 2 major isoforms in immunoglobulin-like domain III (IIIb and IIIc) that influence binding specificity and affinity (Plotnikov et al., 2000; Yeh et al., 2003). Ucp1 expression is induced in preadipocytes by FGF8 through a signaling cascade emanating from FGFR1 and requires both prostaglandin synthesis and a high glycolytic flux (Gantert et al., 2021).

In this study, we screened all paracrine-acting FGFs for their potential to induce Ucp1 expression in preadipocytes, comprehensively describe the resulting cell type, and identified a programming effect of FGFs that is maintained throughout cell differentiation to fine-tune inflammatory status of mature adipocytes.

## Methods

### Mice

All mice were maintained at 23 °C ambient temperature and 55 % relative humidity, with a 12 h light/dark cycle in our specified-pathogen-free mouse facility. Mice had ad libitum access to a standard maintenance and breeding chow diet (Ssniff Spezialdiäten GmbH, Germany) and water. Paracrine FGF screening was performed with primary preadipocytes isolated from the *Ucp1* dual-reporter gene mouse C57BL/6NTac-Ucp1tm3588, simultaneously expressing firefly luciferase and near-infrared fluorescent protein 713 (Wang et al., 2019). Subsequent studies were conducted in primary preadipocytes derived from Bl6N wildtype mice as indicated.

### Cell Culture

All immortalized cells used in this study were cultured and differentiated in high glucose DMEM (4.5 g/L glucose, Sigma-Aldrich) with 10 % FBS (FBS Supreme, PAN Biotech). Primary cells were cultured in 20 % FBS supplemented high glucose DMEM until 80% confluency and treated or differentiated in 10 % FBS supplemented to high glucose DMEM. For treatment of both, primary and immortalized preadipocytes, paracrine FGF recombinant proteins were added in the culture media (R&D Systems). Primary SVF were isolated from iBAT and iWAT of 8-10 week old Bl6N mice. iBAT and iWAT were minced until homogenous and digested using collagenase (Cellsystems). Digested tissues were filtered through a 250 µM nylon mesh. To precipitate SVFs filtered tissues were centrifuged at 250 g for 5 min. The SVF pellet was resuspended in wash buffer (1x HBSS w/Mg;Ca + 3,5% BSA) and centrifuged at 500 g for 5 minutes. Isolated cells were cultured until 80 % confluency before FGF treatment. Subsequent differentiation was induced with an induction cocktail prepared in high glucose DMEM supplemented with 10 % FBS and contained 850 nM insulin, 1 nM T3, 500 μM IBMX, 1 μM dexamethasone, 125 μM indomethacin, and 1 μM rosiglitazone. After 48 hours, the induction medium was replaced with differentiation medium containing 850 nM insulin, 1 nM T3, and 1 μM rosiglitazone. Cells were differentiated for 6 days. FGFR inhibitor (LY2874455, MCE) was used simultaneously at indicated concentrations throughout FGF treatment.

### Transfection of WT-1-CRISPRa-SAM preadipocytes

The WT-1-CRISPRa-SAM cell line is designed to activate the expression of target genes through the CRISPRa Synergistic Activation Mediator (SAM) system (Lundh et al., 2017). The WT-1-CRISPRa-SAM cell line, stably expressing the SAM components, was seeded onto 24-well plates and transfected upon reaching confluency. For transfection, 250 ng of sgRNA targeting either *Ucp1* or *Fgf2* was mixed with TransIT-X2 transfection reagent according to the manufacturer’s instructions and added to each well. Forty-eight hours post-transfection, the cell culture medium was replaced, and after an additional 48 hours, the transfected preadipocytes were harvested or differentiated for subsequent analyses.

### Bioluminescence assay

Luciferase activity was quantified as indirect measure for *Ucp1* expression (*Ucp1*-LUC) in cell lysates derived from the *Ucp1*-reporter mouse model. Luminescence was measured with the InfiniteM2000 NanoQuant, (Tecan) using the luciferase assay system kit (E1501, Promega). Briefly, cells were washed with PBS once and lysed in cell culture lysis reagent for 30 min at 20 rpm. Ten ul of cell lysate was used and added to 50 ul of luciferase assay buffer on a white 96-well plate. Luciferase assay buffer was added and luminescence was measured over an integration time of 10 ms directly after. Luminescence was normalized to non-treated and FGF1-treated cells.

### Respirometry

Immortalized brown preadipocytes were treated with paracrine FGFs (1, 2, 4, 8, and 9) and oxygen consumption rate (OCR) was either directly measured or cells underwent differentiation as stated above following OCR measurement (Oeckl et al., 2020). For preadipocytes, Mir05 was used (supplemented with 25 mM glucose and 5 mM succinate) as assay medium. Fully differentiated adipocytes were measured in Seahorse medium (containing 2% (w/v) essentially fatty-acid-free BSA). A Seahorse XF Pro Sensor Cartridge (Agilent Technologies) was hydrated overnight with XF calibrant and in a CO₂-free incubator at 37 °C. For measurements, cells were washed twice with the appropriate assay medium and 180 μL of assay medium was added to each well. The washed cells were incubated at 37 °C for 1 hour in CO₂-free conditions using the Cytation 1 imaging device (BioTek Cytation 1 Cell Imaging Multimode Reader, Agilent Technologies) to ensure proper outgassing. Bright-field images of the cells were captured automatically during the incubation using Cell Imaging software (Agilent Technologies). To measure Ucp1-dependent respiration in preadipocytes, cells were permeabilized by 20 µM a-chaconine (Dawid et al., 2020). Following cell lysing, 20 µM 4-[(1E)-2-(5,6,7,8-tetrahydro-5,5,8,8-tetramethyl-2-naphthalenyl)-1-propen-1-yl]-benzoic acid (TTNPB) was injected for direct Ucp1 activation. 1 µM carbonylcyanidep-trifluoromethoxy-phenylhydrazone (FCCP) and 5 µM Antimycin A were injected afterwards. Ucp1-dependent respiration in adipocytes was measured with the following injection scheme: 5µM Oligomycin, 0.5 µM Isoproterenol, 1 µM FCCP, 5 µM Antimycin A. Oxygen consumption rate (OCR) was recorded in response to the experimental substrates. Data visualization and initial analysis were performed using Wave Pro software (Agilent Technologies).

### Oil Red O staining

Oil Red O working solution was prepared by mixing three parts of Oil Red O stock solution (0.5 % w/v Oil Red O, 100 % v/v Isopropanol, filtered through 0.45 µM filter) with two parts of double-deionized water (ddH₂O), followed by filtration through a 0.22 μm filter (Berrytec, Cat. No.: 1302601). Fully differentiated cells were washed twice with pre-warmed phosphate-buffered saline (PBS) and fixed with 100 µL/well of 3.7% (v/v) formaldehyde for 1 h at room temperature. Fixed cells were washed three times with ddH₂O and once with 60% isopropanol for 5 minutes, then air-dried. Cells were incubated for 15 minutes in freshly prepared Oil Red O working solution. Excess stain was removed by washing four times with ddH₂O. Stained lipid droplets were visualized under a light microscope at 20x magnification (Axiovert 40 CFL, Carl Zeiss, Cat. No.: 10326743). For quantification and size distribution analysis, images of unstained cells were analyzed using the WimLipid pattern recognition algorithm (Wimasis, Onimagin Technologies).

### Quantitative RT-PCR

Cells were harvested and lysed with TRIsure reagent, followed by RNA extraction using the SV Total RNA Isolation System (Promega). The RNA concentrations were measured using a Nanodrop-1000 instrument. cDNA was synthesized from 500-1,000 ng RNA using the SensiFast cDNA Synthesis Kit (Bioline). RT-qPCR analysis was performed using the LightCycler480 Real-Time PCR System (Roche) using the SensiMix SYBR No-ROX Kit (Bioline). The relative standard curve method, utilizing pooled sample cDNA standards, was employed for quantification. Target gene expression was normalized to the reference genes *Tbp, Hsp90 and Tf2b*. Primer sequences are listen in table S1.

### bulkRNA-sequencing

Primary or immortalized preadipocytes were treated with vehicle or FGF1, FGF2, FGF4, FGF8, or FGF9 for 48 h. FGF-treated cells were either directly harvested or differentiated as stated above and harvested at different time points as indicated. Harvested cells were lyzed in TRIsure and RNA was extracted using the SV Total RNA Isolation System by Promega following the manufacturer’s protocol. RNA quality and integrity were assessed using Agilent Bioanalyzer 2100. High-quality RNA (RIN > 7) was used to prepare libraries using Illumina stranded mRNA Kit, UDI kit. Sequencing was performed on Illumina NovaSeq 6000, generating paired-end reads of 150 bp. Raw reads were trimmed for quality using TrimGalore and aligned to the reference genome, GRCm39, using HISAT2. Gene-level counts were generated with featureCounts and compiled into a count matrix for subsequent analysis. Count data were analyzed using DESeq2 (v1.42.1). Variance-stabilizing transformation (VST) was applied to normalize the data. PCA and heatmaps were visualized with ggplot2 (v. 3.5.2.) and pheatmap. All analyses were conducted in R (v. 4.3.3).

### Single Cell RNA-sequencing

Primary preadipocytes isolated from iBAT and iWAT were cultured until 80% confluency and subsequently treated with vehicle, FGF1 or FGF2 as stated above for 48 h. Cell suspensions were loaded into the BD Scanner™ to count cells and assess cell viability by brightfield and dual band fluorescence imaging. Samples where tagged using Sample Tags from BD and merged. In total, 40,000 cells were loaded into the BD Rhapsody Cartridge™. Here, one cell were paired with one barcoded bead. Cells were lysed and mRNA was hybridized to the bead. mRNA attached to beads were retrieved magnetically. cDNA synthesis and library preparation was prepared as stated by the supplier. Samples were sequenced on a NovaSeq X Plus (PE150), Illumina. Reads were aligned and annotated using the data pipeline at Seven Bridges Genomics. Subsequent analysis was performed using Seurat (v. 5.0.0). Mitochondrial gene expression percentages were calculated for each cell to assess overall cell quality. Cells were filtered based on the number of detected features per cell (nFeature_RNA > 200 and < 7500) and the percentage of mitochondrial transcripts (<30%). Normalization of the dataset was performed using log-normalization to adjust for variability in sequencing depth. To account for confounding effects of cell-cycle variation, cell-cycle phase scores were computed using previously published gene markers (Kowalczyk et al., 2015). A difference score was calculated (S.Score - G2M.Score) and regressed out during data scaling. Dimensionality reduction was performed using principal component analysis (PCA), and the optimal number of dimensions for clustering was determined by Elbow Plot. Clustering was conducted using shared nearest neighbor (SNN) graphs, and the clusters were visualized using uniform manifold approximation and projection (UMAP). Clusters were annotated to cell subtypes by known marker genes based on the marker genes for each cluster. Known cell type marker genes were collected from CellMarker2.0 (Hu et al., 2023).

### Statistical analysis

Statistical analyses were conducted using GraphPad Prism version 10.4.2 (GraphPad Software, San Diego, California, USA). Parametric tests, including Student’s t-test and one-way or two-way ANOVA with Bonferroni correction, were applied to data that followed an approximately normal distribution. Data are reported as means ± SD unless specified otherwise in the figure legends. Statistical significance was set at p < 0.05 and denoted with asterisks: *p < 0.05.

## Results

### 1 Paracrine fibroblast growth factors 2, 4, 8, and 9 induce Ucp1 protein expression in primary white and brown preadipocytes

Fibroblast growth factor 8 (FGF8) induces the atypical expression of uncoupling protein 1 (Ucp1) in white and brown preadipocytes via FGF receptor 1 (FGFR1) (Gantert et al., 2021). The FGFR1 is a promiscuous receptor interacting with several FGFs with varying affinity (Ornitz et al., 1996). We conducted a comprehensive screening using primary preadipocytes derived from a luciferase-based Ucp1 reporter mouse to explore the potential of all paracrine acting fibroblast growth factors (FGFs 1-10, 16-18, 20, and 22) to drive Ucp1 expression. The model features a luciferase open reading frame inserted into the endogenous Ucp1 locus, enabling indirect measurement of Ucp1 gene transcriptional activity via luminescence (Wang et al., 2019). Primary preadipocytes from both interscapular brown adipose tissue (iBAT) and inguinal white adipose tissue (iWAT) were harvested and treated with each of the FGFs, respectively. The factors FGF2, 4, 8 and 9 proved to be potent inducers of Ucp1 expression across primary preadipocytes derived from both iBAT and iWAT, with FGF2 eliciting the strongest response, followed by FGF8, FGF9, and FGF4 (Fig. 1 A, B). The highest level of Ucp1 expression was reached after a 48-hour treatment (Fig. 1 C). In dose-response experiments, FGF2 and 8 induced Ucp1 reporter activity at lowest (nanomolar) concentrations (Fig. 1 D-G). The FGFs 4 and 9 required higher concentrations to initiate Ucp1 reporter activity, and no plateau was yet reached at the maximum tested concentration of 100 nM.

**Figure 1.**
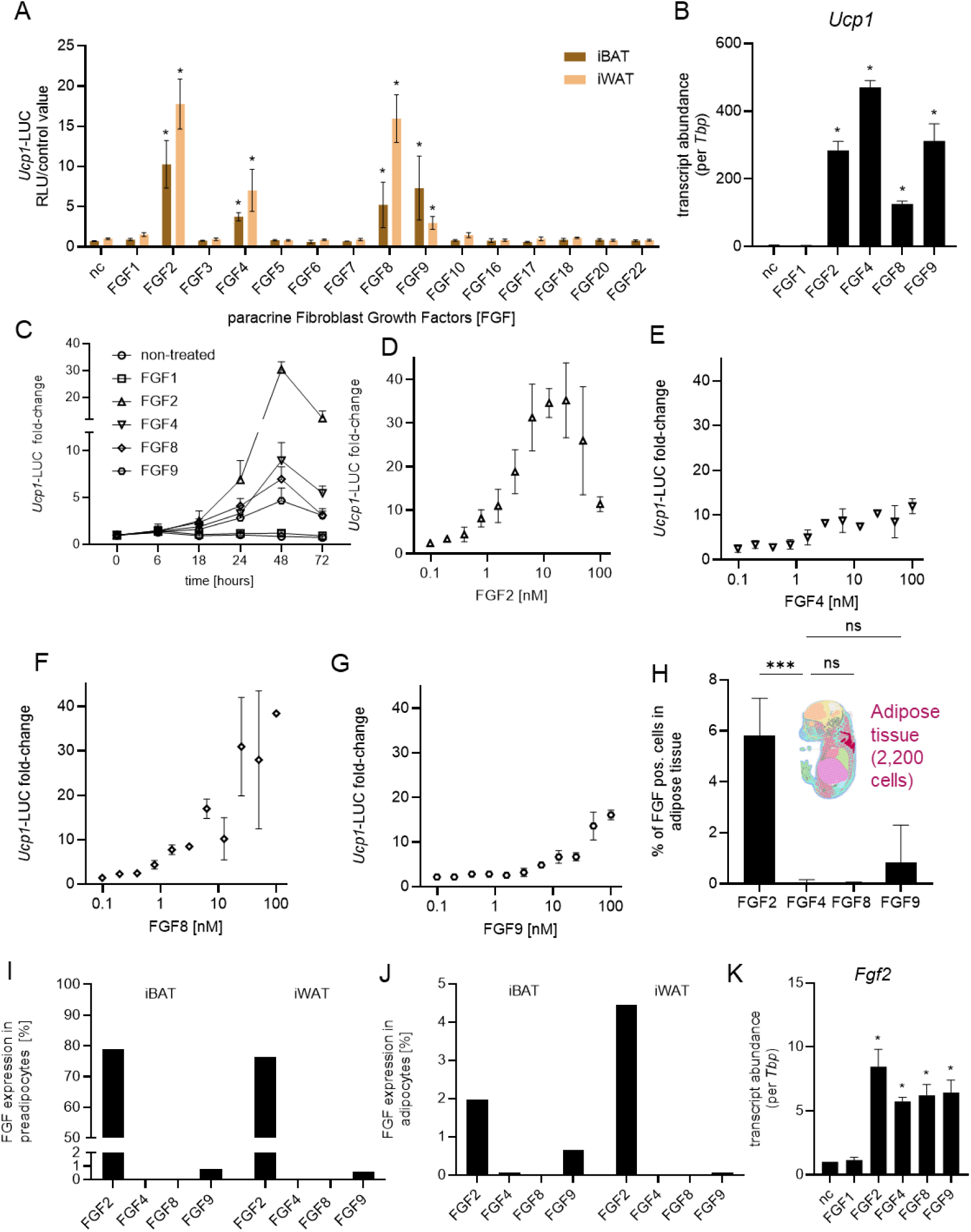
Paracrine fibroblast growth factors 2, 4, 8, and 9 induce Ucp1 expression in brown and white preadipocytes. **(A)** Luciferase UCP1 reporter activity upon treatment with individual FGFs in brown and white preadipocytes from *Ucp1*-reporter mice. Data represent mean ± SD; *n* = 3. Statistical analysis: two-way ANOVA with Dunnett’s multiple comparisons test. **(B)** *Ucp1* mRNA abundance normalized to *Tbp* in immortalized brown preadipocytes treated with 5.5 nM FGF1, FGF2, FGF4, FGF8, or FGF9 for 48 hours, compared to untreated controls. Data represent mean ± SD; *n* = 3. Statistical analysis: one-way ANOVA with Dunnett’s test. **(C)** Time-course of *Ucp1*-linked luminescence in white preadipocytes treated with 5.5 nM FGF1, FGF2, FGF4, FGF8, or FGF9. Data represent mean ± SD; *n* = 3 biological replicates. Statistical analysis: two-way ANOVA with Dunnett’s multiple comparisons test. **(D–G)** Dose–response curves of *Ucp1* reporter luminescence in white preadipocytes treated with increasing concentrations of FGF1, FGF2, FGF4, FGF8, or FGF9 (maximum 100 nM, 10 serial two-fold dilutions). Data represent mean ± SD; *n* = 3 biological replicates. **(H)** Expression of FGF2, FGF4, FGF8, and FGF9 in annotated embryonic brown adipose tissue (E16.5) extracted from the spatial transcriptome database MOSTA (Chen et al., 2022). Data represent mean ± SD; n = 3 biological replicates. Statistical analysis: one-way ANOVA with Tukey’s multiple comparisons test. **(I)** Expression of FGF2, FGF4, FGF8, and FGF9 in preadipocytes from adult iWAT and iBAT, extracted from our single-cell RNA-seq dataset. Data represent one dataset per tissue. **(J)** Expression of FGF2, FGF4, FGF8, and FGF9 in mature adipocytes from adult iWAT and iBAT, based on single-nucleus RNA-seq data. Data represent one dataset per tissue. **(K)** *Fgf2* mRNA abundance normalized to *Tbp* in immortalized brown preadipocytes treated with 5.5 nM FGF1, FGF2, FGF4, FGF8, or FGF9 for 48 hours, compared to untreated controls. Data represent mean ± SD; *n* = 3. Statistical analysis: one-way ANOVA with Dunnett’s test. In all panels: * = p<0.05.

FGFs are recognized primarily for their roles in embryonic development, particularly in brain, limb, and central nervous system formation (Ornitz & Itoh, 2022). Both white and brown adipose tissue depots develop late during embryogenesis (Seale et al., 2008) and FGFs may play a physiological role during these events. We thus explored the presence of Ucp1-inducing FGFs 2, 4, 8, and 9 in both mature and developing adipose tissues. We re-analyzed the publicly available Mouse Organogenesis Spatiotemporal Transcriptomic Atlas (MOSTA, (Chen et al., 2022)) as well as own datasets, i.e. single-cell RNA-sequencing (scRNA-seq) datasets from brown and white primary preadipocytes, and single-nuclei RNA-sequencing (snRNAseq) datasets from mature brown and white adipocytes.

In developing adipose tissue, FGF2 displayed the highest expression compared to FGF4, 8, and 9 at embryogenic stage E16.5 (Fig. 1 H). In primary preadipocytes from both inguinal white and interscapular brown adipose tissue, only FGF2 was consistently expressed, while FGF4, FGF8, and FGF9 were absent in preadipocyte populations (Fig. 1 I). Similarly, FGF2 was the only present FGF in mature adipocytes derived from iWAT and iBAT (Fig. 1 J). Taken together, these findings point towards FGF2 as a relevant driver of Ucp1 expression in vivo. Interestingly, treatment with FGF4, FGF8, or FGF9 (but not control FGF1) consistently increased FGF2 expression in preadipocytes, supporting FGF2 as the primary mediator of FGF-induced Ucp1 expression (Fig. 1 K).

In summary, FGFs 2, 4, 8, and 9 act as inducers of significant Ucp1 transcript expression in both inguinal white and interscapular brown preadipocytes. Of these, FGF2 is predominantly expressed at all developmental stages of adipose tissue in mice as well as in mature adipose tissue preadipocytes and adipocytes.

### 2 Fibroblast growth factors 2, 4, 8, and 9 signal via the same pathway to induce Ucp1 but do not induce a ‘browning’ signature

The FGFR1 is essential for the induction of Ucp1 expression by FGF8 (Gantert et al., 2021). The binding affinities of FGF2, 4, 8, and 9 to the group of FGF receptors are well in line with a common signaling path via FGFR1 (Ornitz & Itoh, 2022). FGF2 binds most strongly to the receptor isoforms FGFR1b and FGFR1c; FGF4 primarily binds FGFR1c and FGFR2c; FGF8 has highest affinity for FGFR1c; and FGF9 binds most strongly to FGFR1c. Thus, FGFR1 isoforms are the likely receptors mediating the FGF-Ucp1 axis in preadipocytes for all identified FGFs, even in the case that FGFs 4, 8, and 9 would not act via induction of the primary active agent FGF2.

We characterized the transcriptional response to various FGFs by RNA sequencing of inguinal white primary preadipocytes treated with FGFs. Principal Component Analysis (PCA) of FGF-treated cells revealed a distinct effect of treatment compared to non-treated and FGF1-treated cells (Fig. 2 A). Clustering along the first principal component reflected the strength of Ucp1 induction across FGFs, with FGF2 as the strongest inducer and FGF4 the weakest. FGFs 2, 4, 8 and 9 did not form separate clusters along any other principal component, corroborating a shared receptor and signaling pathway. Similarly, in a heatmap analysis of the most FGF-sensitive transcripts, non-treated and FGF1-treated cells displayed a similar gene expression profile different from a shared pattern induced by FGF2, FGF4, FGF8, and FGF9 (Fig. 2 B). Marker transcripts of adipocyte browning or differentiation were, however, not FGF-induced (Fig. 2 C). Surprisingly, the induction of the hallmark brown/beige adipocyte marker Ucp1 was not part of an accompanying generalized browning phenomenon but occurred separate from the well-characterized thermogenic expression signature.

**Figure 2.**
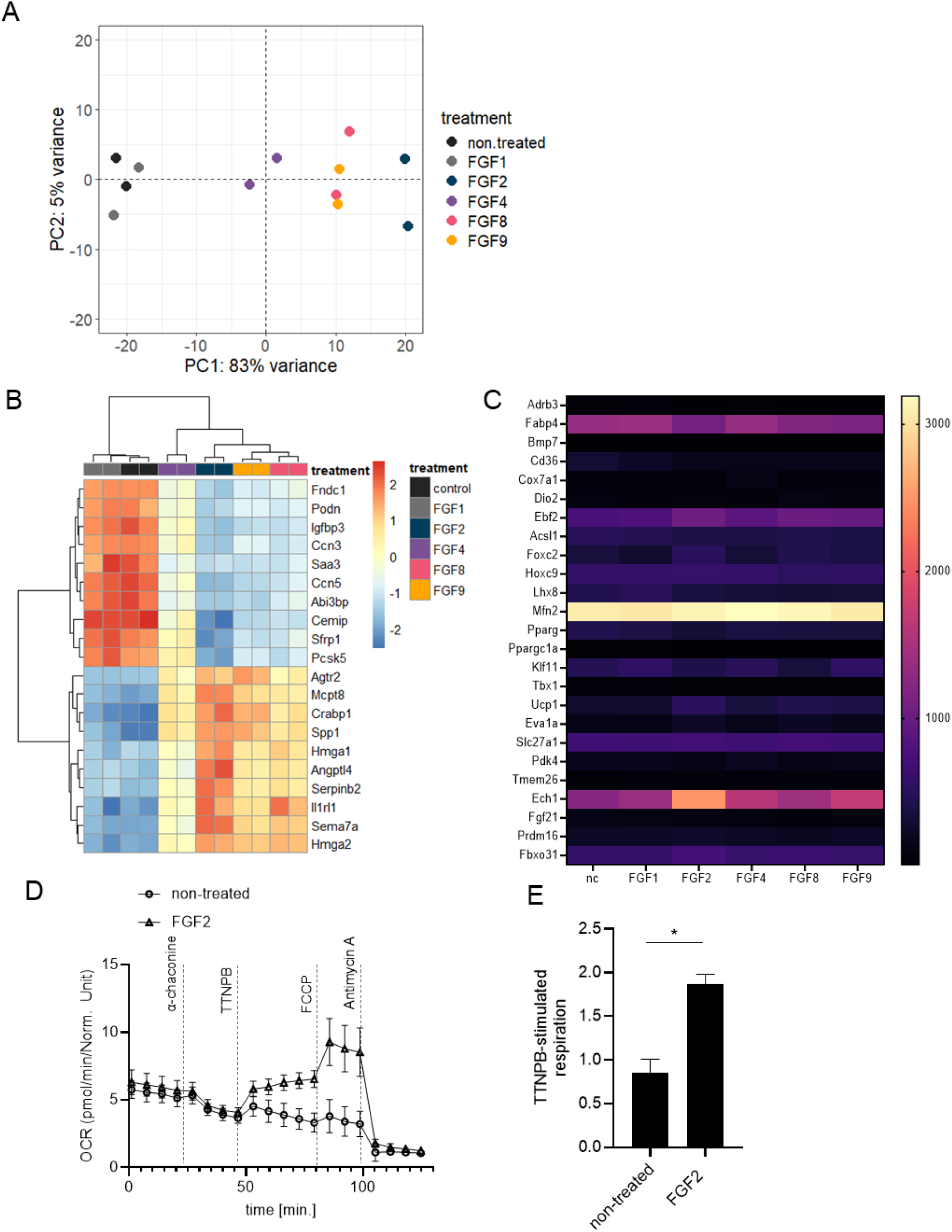
FGF2, FGF4, FGF8, and FGF9 induce a shared transcriptomic signature in white preadipocytes. **(A)** Principal component analysis (PCA) of global transcriptomic profiles from white preadipocytes treated with FGF1, FGF2, FGF4, FGF8, or FGF9, compared to untreated controls. Each condition includes *N* = 2 biological replicates. PCA was performed using variance-stabilized transformed counts. **(B)** Heatmap showing the top 20 most variable genes across all samples, selected based on variance-stabilized expression values. Data are clustered by gene and sample to illustrate global transcriptional differences between treatments. **(C)** Heatmap displaying the expression of 25 selected browning-associated genes in white preadipocytes treated with 5.5 nM FGF2, FGF4, FGF8, or FGF9 for 48 hours, compared to untreated controls. Gene expression is based on normalized counts from bulk RNA-seq data. Rows represent genes, and columns represent treatment groups. **(D)** Time course of oxygen consumption rate in immortalized brown preadipocytes comparing FGF2 treatment with its untreated control, measured by microplate-based respirometry (Seahorse XF96 Analyzer). Oxygen consumption is recorded in lyzed cells under basal conditions and in response to successive injection TTNPB (1 µM), FCCP (1 µM), and antimycin A (5 µM) to determine direct UCP1 activity, maximal and non-mitochondrial respiration, respectively. Mean values ± SD, n = 3 (biological replicates). **(E)** Quantification of TTNPB-stimulated UCP1-mediated uncoupled respiration, expressed as fold of basal leak respiration. Mean values ± SD, n = 3 (biological replicates). Statistical analysis: unpaired t-test.

The absence of a typical environment that would enable thermogenic Ucp1 activity in differentiated brown adipocytes prompted us to determine the functionality of Ucp1. Since preadipocytes lack beta-adrenergic receptors and relevant intracellular lipid stores — both required to activate Ucp1-mediated mitochondrial uncoupling — we utilized a direct activator of Ucp1 activity, the retinoic acid analog 4-[(1E)-2-(5,6,7,8-tetrahydro-5,5,8,8-tetramethyl-2-naphthalenyl)-1-propen-1-yl]-benzoic acid (TTNPB) (Rial, 1999). Remarkably, Ucp1 in FGF2-treated preadipocytes could indeed be activated to a considerable portion of maximal oxidative capacity, demonstrating proper location to the inner mitochondrial membrane and principal functionality (Fig. 2 D, E).

In summary, all four paracrine FGFs 2, 4, 8, and 9 induced the same transcriptomic signature in primary white preadipocytes indicating a common receptor and signaling pathway. They do not initiate browning *per se* or differentiation. Importantly, Ucp1 protein derived from FGF2 treatment was functional, suggesting an alternative role for Ucp1 in this cell type different from *bona fide* thermogenesis.

### 3 Single-cell analysis of FGF 2 - driven preadipocyte transition reveals persisting preadipocyte identity of Ucp1 - expressing cells

The cellular identity and biological role of FGF-induced, Ucp1-expressing cells is unclear. We performed single-cell RNA sequencing (scRNA-seq) to analyze which primary cell population is FGF2-responsive and if such treatment induces a new and discrete Ucp1-positive subpopulation. Uniform Manifold Approximation and Projection (UMAP) dimensionality reduction of primary cells isolated and cultured from white and brown adipose tissue revealed 13 distinct clusters in iWAT and 12 clusters in iBAT, representing transcriptionally distinct cell populations (Fig. 3 A, B). These were manually annotated by marker genes derived from the CellMarker 2.0 database (Hu et al., 2023). Three prominent clusters corresponding to preadipocytes and fibroblast-like cell populations were identified along with smaller clusters representing immune and endothelial cells. In iWAT-derived cells, FGF2 treatment caused a shift in subpopulations within the pre-existing transcriptional space of identified preadipocytes (Fig. 3 C, D). Similarly, in iBAT-derived cells, FGF2 treatment induced a pronounced shift in cell distribution within preadipocyte clusters (Fig. 3 E, F). Importantly, no entirely new subpopulation of preadipocytes or any other distinct cell type emerged in response to FGF2 treatment (Fig. 3 D, F). Thus, neither in primary cells from iWAT nor iBAT FGF2 re-programmed preadipocytes into an entirely new subpopulation but rather altered the distribution into pre-existing preadipocyte sub-populations.

**Figure 3.**
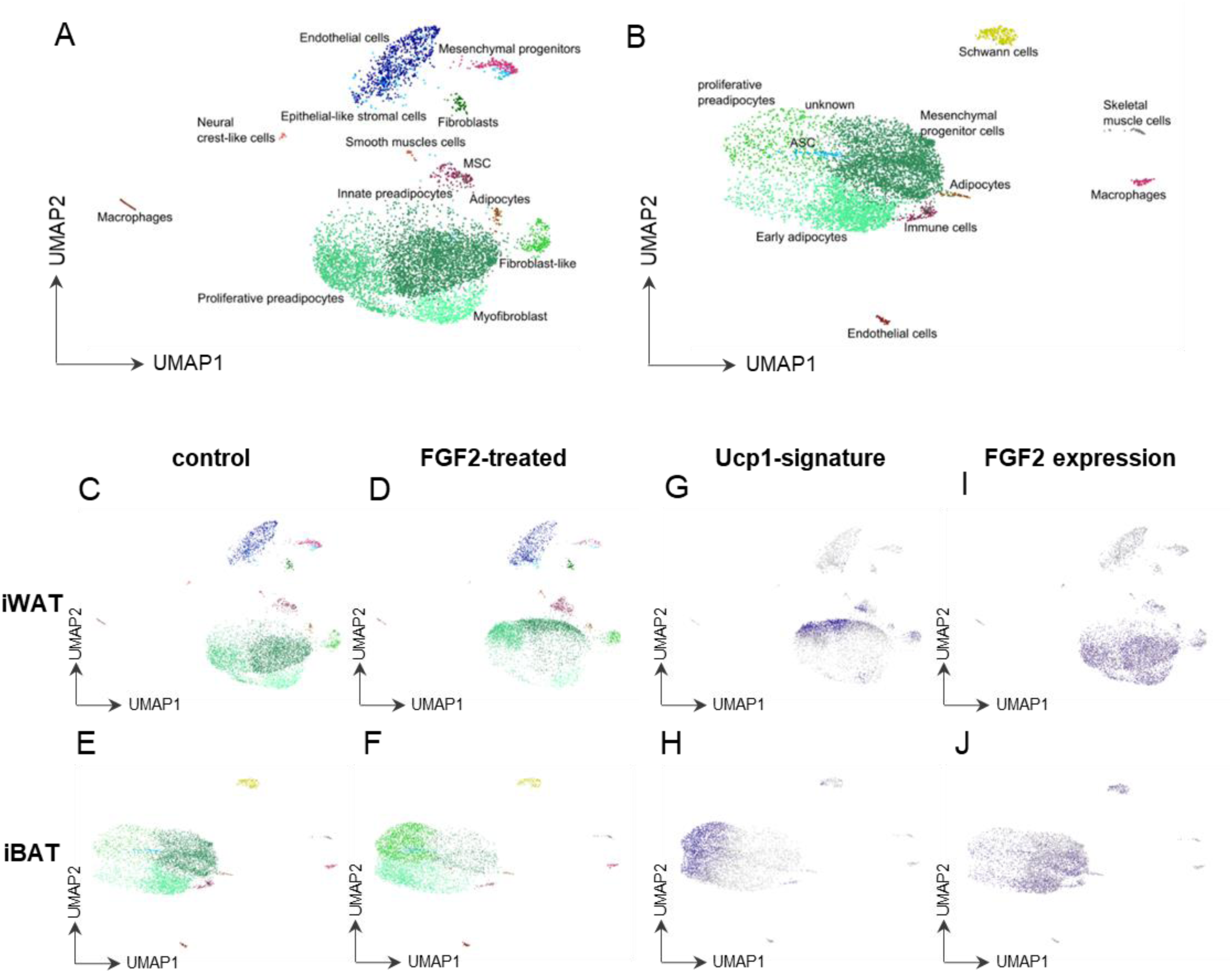
FGF2 treatment reshapes the cellular landscape of primary iWAT and iBAT preadipocytes. **(A)** UMAP of iWAT-derived cells (stromal-vascular fraction), revealing 13 distinct clusters. In the untreated condition, preadipocytes constitute the predominant population. **(B)** UMAP of iBAT-derived cells(stromal-vascular fraction), showing 12 distinct clusters. Similar to iWAT, untreated samples are primarily composed of preadipocytes. **(C)** UMAP of untreated iWAT preadipocytes showing the baseline distribution of clusters and **(D)** UMAP of FGF2-treated iWAT preadipocytes reveals a shift in cluster composition within the preadipocyte population. **(E)** UMAP of untreated iBAT preadipocytes showing the baseline cluster distribution and **(F)** UMAP of FGF2-treated iBAT preadipocytes displaying a similar shift in cluster localization within the preadipocyte compartment. **(G)** Expression of a Ucp1 gene signature (top 20 upregulated genes from bulk RNA-seq) in FGF2-treated iWAT preadipocytes, specifically localized to the shifted preadipocyte population and **(H)** Expression of the Ucp1 gene signature in FGF2-treated iBAT preadipocytes, marking the same preadipocyte subpopulation that shifts upon treatment. **(I)** FeaturePlot showing endogenous FGF2 expression in untreated iWAT preadipocytes **(J)** and in untreated iBAT preadipocytes. Cells expressing *Fgf2* are highlighted in blue; non-expressing cells are shown in grey with *Fgf2* expression restricted to the subset of cells previously shown to respond with a *Ucp1*-associated transcriptional shift.

We verified the FGF2-induced shift in preadipocyte characteristics to be associated with the initially observed induction of Ucp1 expression. The top 20 transcripts correlated to Ucp1 expression in the previous bulk RNA sequencing experiment (Fig. 2 B) mapped very well onto the preadipocyte population arising in response to FGF2 treatment (Fig. 3 G, H). This Ucp1 associated signature was observed in primary preadipocytes from both iBAT and iWAT demonstrating that both respond to FGF2 by positively regulating Ucp1 without forming a new, non-preadipocyte subpopulation.

The signal FGF2 itself was already expressed by cell populations within the untreated primary cells, indicating ongoing FGF2 crosstalk in the absence of any manipulation (Fig. 3 I, J). Interestingly, these FGF2 positive cells themselves were nearly exclusively of the preadipocyte group, indicating a paracrine and/or autocrine path of communication.

Pathway analyses of FGF2-responding preadipocytes demonstrated a significant upregulation in cell proliferation pathways, consistent with prior findings of FGF8-mediated effects (Gantert et al., 2021). Pathways related to browning or adipogenesis were not enriched, in line with our earlier interpretation of global transcriptome changes. Ucp1 expression in preadipocytes is independent of these well-known differentiation processes and thus represents a distinct phenomenon within the preadipocyte lineage.

In summary, single-cell RNA sequencing of iWAT and iBAT preadipocytes identified a distinct, naturally occurring preadipocyte population responsive to FGF2 treatment. Responding preadipocytes remained within the bounds of pre-existing preadipocyte clusters rather than forming a new population. FGF2 itself also localized to preadipocytes, suggesting an autocrine and/or paracrine signaling path within the FGF2-Ucp1 axis. Pathway analyses confirmed the absence of browning or adipogenesis induction, highlighting a unique FGF2-driven Ucp1 expression in preadipocytes while preserving their overall cellular identity.

### 4 Fibroblast growth factor-pretreated adipocytes differentiate into fully functional adipocytes

Early exposure to FGF2, FGF4, FGF8, or FGF9 induces Ucp1 expression in a resulting cell population that otherwise retains the identity of preadipocytes. We tested the hypothesis that fully differentiated adipocytes originating from these cells differ from adipocytes differentiated under control condition. After FGF pre-treatment, we induced differentiation using standard induction and differentiation cocktails (Fig. 4 A). After seven days of differentiation (i.e. nine days since the end of FGF treatment), resulting adipocytes appeared morphologically indistinguishable upon visual inspection, FGF-treated or not (Fig. 4 B). Likewise, fat accumulation into lipid droplets remained unchanged, both measured as total fat content and as lipid droplet number and size distribution (Fig. 4 C, D). In line, thermogenic capacity of Ucp1 in response to isoproterenol stimulation remained similar in control and FGF-pretreated brown adipocytes (Fig. 4 E, F). Taken together, adipocytes differentiated from FGF-treated preadipocytes are fully functional and morphologically unchanged compared to untreated cells. The FGF-induced, Ucp1-positive preadipocyte population does not give rise to a different functional class of mature adipocyte.

**Figure 4.**
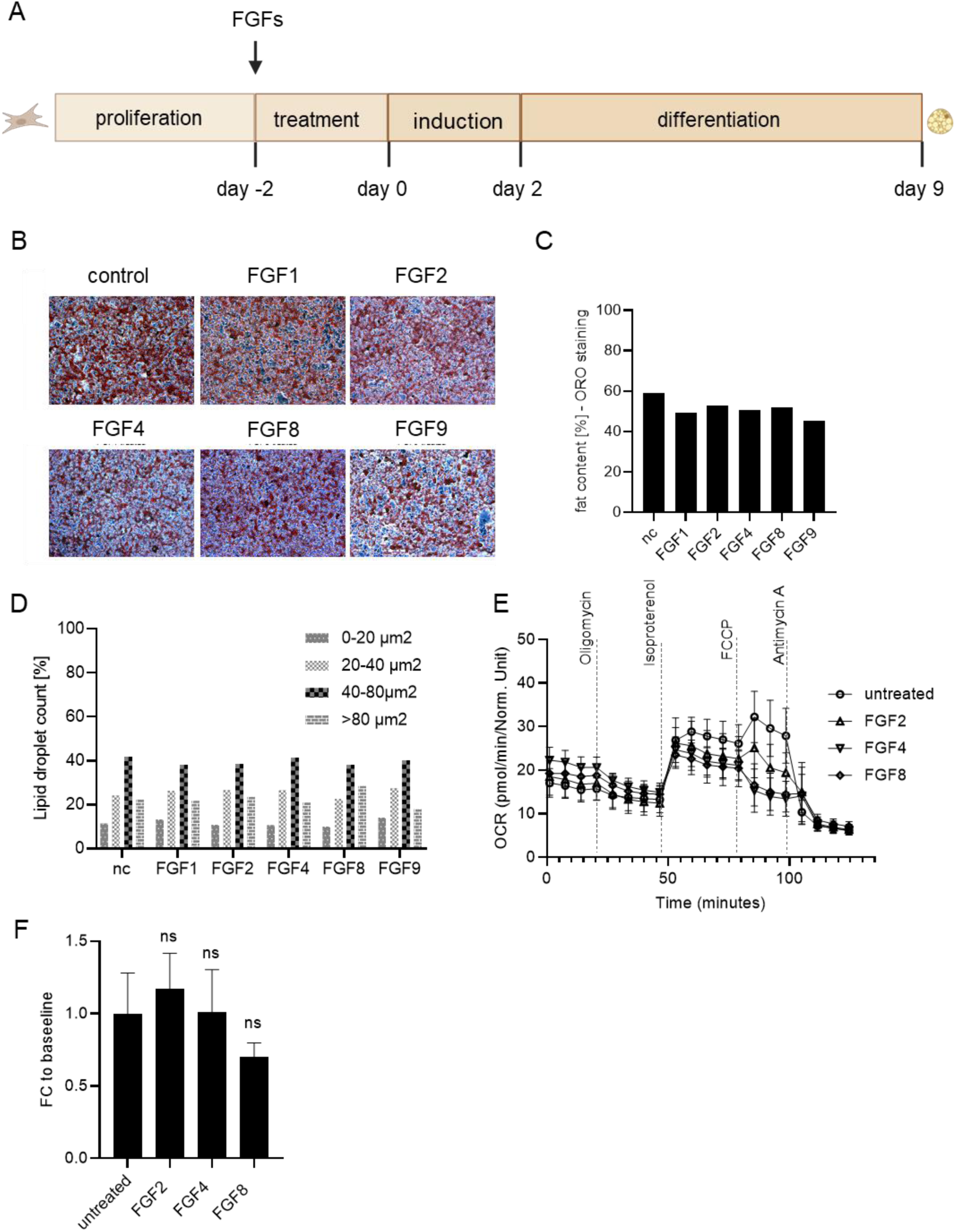
FGF pretreatment does not alter adipocyte lipid accumulation or droplet morphology when differentiated into adipocytes. **(A)** Experimental design showing preadipocyte treatment with FGF1, FGF2, FGF4, FGF8, or FGF9 followed by differentiation. **(B)** Oil Red O staining of lipid droplets in differentiated adipocytes pretreated with individual FGFs, compared to untreated controls. **(C)** Quantification of lipid content as percentage of total well area stained, showing no significant differences across treatments. **(D)** Lipid droplet count per treatment group, stratified by droplet size (0–20 µm², 20–40 µm², 40–80 µm², >80 µm²), revealing similar distributions across all conditions. **(E)** Time course of oxygen consumption rate in immortalized brown adipocytes pretreated with FGF2, 4, or 8 comparing with its untreated control, measured by microplate-based respirometry (Seahorse XF96 Analyzer). Oxygen consumption is recorded under basal conditions and in response to successive injection of oligomycin (5 µM), isoproterenol (1 µM), FCCP (1 µM), and antimycin A (5 µM) to determine basal leak, UCP1-dependent uncoupled, maximal and non-mitochondrial respiration, respectively. Mean values ± SD, n = 3 (biological replicates). **(F)** Quantification of isoproterenol-stimulated UCP1-mediated uncoupled respiration, expressed as fold of basal leak respiration. Mean values ± SD, n = 3 (biological replicates). Statistical analysis: paired t-tests.

### 5 Fibroblast growth factor induced preadipocyte Ucp1 expression causes lasting changes to the interferon pathway in mature adipocytes

Mature adipocytes differentiated from FGF-treated, Ucp1-positive preadipocytes appeared morphologically and functionally unaltered. We conducted a global transcriptome analysis of FGF-treated preadipocytes and adipocytes at day 3 and 7 of differentiation to detect other consequences of FGF-pretreatment. After acute treatment, we again observed Ucp1 induction and the associated expression signature in preadipocytes, consistent with previous findings (Fig. 5 A). Over the course of differentiation, Ucp1 induction gradually diminished to the level of untreated adipocytes. On day 7 of differentiation, none of the acutely upregulated transcripts remained different from controls. However, a principal component analysis (PCA) of non-treated and FGF pre-treated preadipocytes and adipocytes revealed a distinct and persistent separation of pre-treated and non-treated cells at all three differentiation states, proving a lasting effect of FGF treatment on the transcriptional profile. This signature was dominated by down-, not up-, regulated transcripts (Fig. 5 B). We applied a pathway analysis to identify the underlying pattern and detected significant attenuation of several pathways associated with interferon and immune response signaling (Fig. 5 C). Specifically, a large set of well-known interferon beta downstream targets remained consistently and strongly reduced across all differentiation states, among them the key players of adipose tissue inflammation, interleukin-6 (IL6) and tumor necrosis factor (Tnf) (Fig. 5 D).

**Figure 5.**
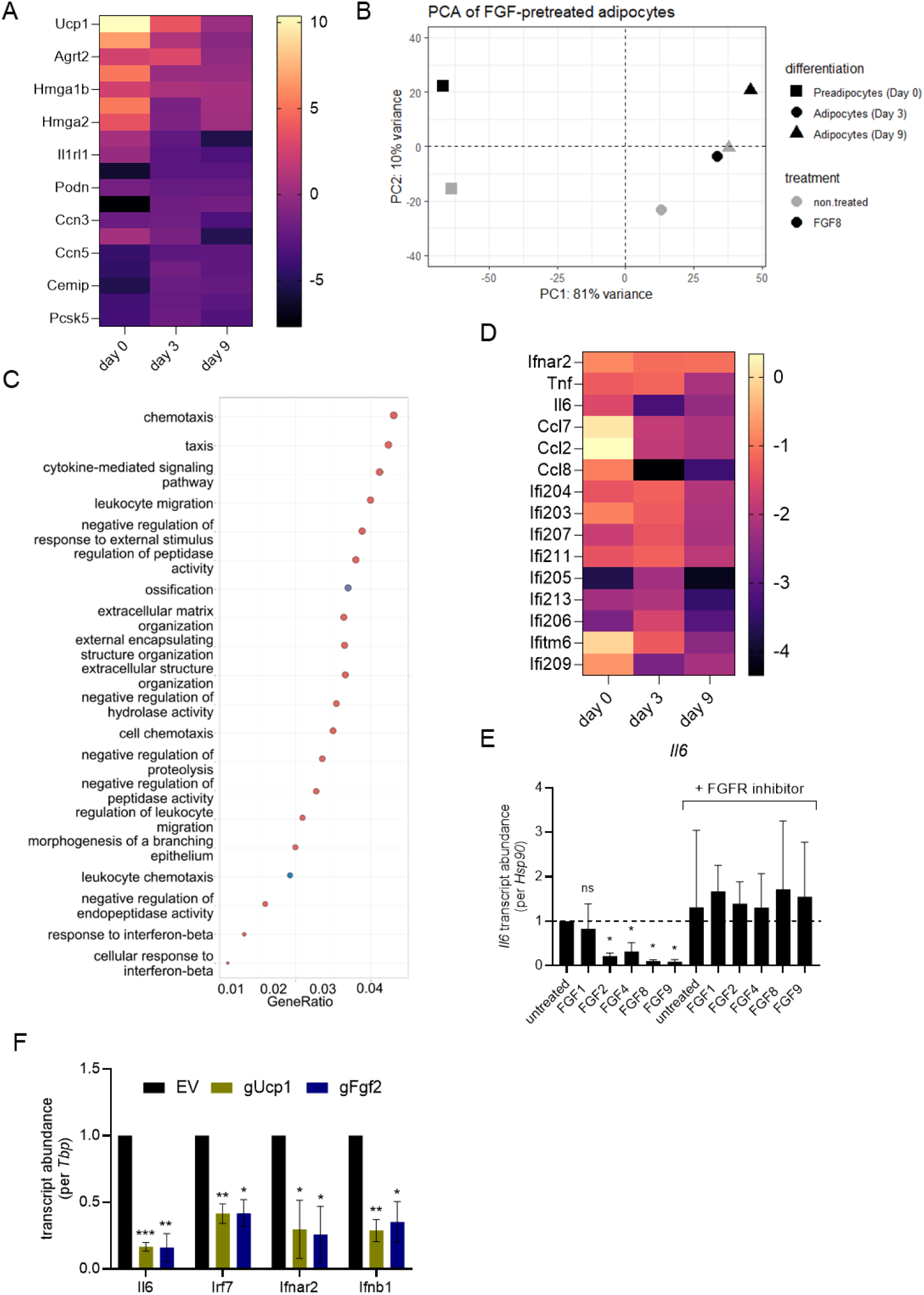
UCP1 mediates FGF-induced suppression of inflammatory gene expression in brown adipocytes. **(A)** Heatmap showing fold changes in expression of Ucp1 signature genes—initially upregulated by FGF treatment in preadipocytes— across preadipocyte, day 3, and day 9 stages, relative to untreated controls. **(B)** Principal component analysis (PCA) of transcriptomic profiles from brown preadipocytes pretreated with FGF8 followed by differentiation and bulk RNA sequencing at early (day 3) and late (day 9) stages. PCA was performed on variance-stabilized transformed counts. **(C)** Gene Ontology (GO) term enrichment analysis of differentially expressed genes reveals suppression of cytokine-mediated signaling pathways in FGF8-pretreated adipocytes. **(D)** Heatmap of inflammatory genes across preadipocyte, day 3, and day 9 time points, displaying fold change relative to untreated controls. Inflammatory gene expression remains broadly reduced following FGF8 pretreatment. **(E)** *IL6* expression in primary brown adipocytes is reduced by FGF2, 4, 8, and 9 pretreatment; this effect is abolished by FGFR inhibition. Data represent qPCR from differentiated cells after FGF pretreatment with or without FGFR inhibitor. Data represent mean ± SD; *n* = 3. Statistical analysis to compare FGF treated cells: one-way ANOVA with Dunnett’s test. **(F)** Overexpression of *Ucp1* or *Fgf2* via CRISPRa in brown preadipocytes suppresses *IL6*, *Irf7*, *Ifnar2*, and *Ifnb1* expression after differentiation, showing that *Ucp1* alone is sufficient to mediate the anti-inflammatory response.

We corroborated this initial finding by treating primary iBAT-derived preadipocytes with FGF1, FGF2, FGF4, FGF8, or FGF9, followed by differentiation. In mature adipocytes, FGF2, FGF4, FGF8, and FGF9 pretreatment strongly reduced *IL6* expression, while FGF1 did not (Fig. 5 E), reproducing our earlier observation (Fig. 5 D). This decrease in *IL6* did not occur in the presence of a pan-FGF receptor inhibitor (during preadipocyte FGF treatment), demonstrating a specific FGFR mediated action of these FGFs. Notably, every single mean of the six conditions with FGFR inhibition displayed a higher *IL6* transcript abundance compared to both control conditions (untreated and FGF1 w/o FGFR inhibitor). Thus, the endogenous FGF2 crosstalk between preadipocytes detected earlier (Fig. 3 I, J) imprints on future *IL6* transcriptional activity nine days later after full differentiation to a mature adipocyte (Fig. 5 E).

The suppression of inflammatory gene expression in FGF8-pretreated adipocytes raised the question of whether the presence of Ucp1 in *preadipocytes* functionally contributes to this state in *mature adipocytes*. Indeed, brown adipose tissue of Ucp1-KO mice has been reported to feature an inflammatory phenotype including high levels of IL6 (Kazak et al., 2017). The putative requirement of early Ucp1 in preadipocytes for later proper control of adipocyte *IL6* expression prompted us to explore a possible permissive role of Ucp1. We utilized a model of CRISPRa-mediated overexpression of *Fgf2* or *Ucp1* in preadipocytes. Overexpression of *Fgf2* in preadipocytes leads to FGF2 release and paracrine/autocrine signaling mimicking exogenous FGF2 treatment including the induction of Ucp1, as demonstrated earlier (Gantert et al., 2021). Intriguingly, forced expression of *Ucp1* in preadipocytes led to a similar downregulation of *IL6* after full differentiation compared to forced *Fgf2* expression (Fig. 5 F). We extended the analysis to other transcripts previously identified to be negatively pre-programmed by FGF2 pre-treatment (Fig. 5 D), namely beta interferon (Ifnb1), its receptor component interferon beta receptor beta chain (Ifnar2) and prominent downstream target interferon regulatory factor 7 (Irf7). All of these were also strongly reduced in mature adipocyte transcript abundance by both early FGF2 and early Ucp1 expression in preadipocytes. Thus, Ucp1 proved necessary and sufficient for FGF2-driven programming of transcriptional activity of the *IL6* and other inflammatory genes, linking the presence of Ucp1 in preadipocytes to the inflammatory status of mature adipocytes.

## Discussion

Fibroblast growth factors (FGFs) are a diverse family of proteins involved in a plethora of developmental and metabolic processes. The subset of paracrine FGFs is characterized by the presence of a heparin binding domain that both limits mobility in the extracellular space and serves as a stabilizer during receptor interaction (Ornitz & Itoh, 2022). We and others previously reported the ability of several paracrine FGFs to induce the unusual presence of UCP1 protein in preadipocytes, prior to initiation of adipogenic differentiation (Gantert et al., 2021; Shamsi et al., 2020; Westphal et al., 2019). In this study, we systematically compared all paracrine FGFs for this ability and identified four FGFs—specifically FGF2, FGF4, FGF8, and FGF9— to induce UCP1 expression in both murine iBAT and iWAT preadipocytes. Among these, FGF2 elicited the strongest and most rapid *Ucp1* induction, was the only FGF endogenously present in appreciable amounts in preadipocytes and developing adipose tissue, and was secondarily induced by the other three candidate FGFs. These arguments render FGF2 the most likely physiological FGF acting on preadipocytes in vivo, in line with its capacity to trigger especially sustained FGFR signaling (Koledova et al., 2019).

In principle, we found UCP1 protein to be functional when activated by the directly interacting activator TTNPB. The well-established canonical signaling cascade activating UCP1 mediated thermogenesis in brown adipocytes, however, is unlikely to be functional in preadipocytes lacking lipid droplets and adrenergic receptors. Furthermore, FGF-induced UCP1 expression in preadipocytes was not accompanied by the well-established gene expression signature enabling thermogenesis in brown and beige adipocytes. Indeed, FGF2/4/8/9 treatment did not give rise to an entirely novel cell population despite the untypical presence of UCP1. Instead, UCP1-positive cells retained their preadipocyte identity and joined a previously rare, but pre-existing preadipocyte subpopulation. Under control conditions, this population is likely the product of endogenous, paracrine/autocrine FGF2 signaling, as detected specifically among the preadipocytes of stromal-vascular fraction.

Full differentiation of FGF-treated, UCP1-positive preadipocytes did not lead to increased Ucp1 abundance or activity in resulting mature brown adipocytes, nor did it induce morphological or adipogenic changes. It did, however, manifest in a strongly reduced inflammatory gene expression signature, specifically of targets of beta interferon, including the hallmark cytokine of adipose tissue inflammation, IL6 (Han et al., 2020). Beta interferon has been described as master regulator of adipose tissue inflammation (Chan et al., 2020). Both beta interferon and IL6 serve as vital hubs of mitogenic and thermogenic control in adipocytes (Kissig et al., 2017; Radványi & Röszer, 2024). Notably, these changes were detected in mature adipocytes, nine days after cessation of FGF treatment and after full adipogenic differentiation, demonstrating a highly persistent imprint on transcriptional control. This phenomenon also occurred endogenously in a crosstalk of primary preadipocytes, as demonstrated by an opposing imprint caused by FGF receptor inhibition.

Intriguingly, the initial finding of atypical UCP1 expression proved mechanistically linked to re-programmed IL6 and other inflammatory markers in resulting adipocytes. In BAT of UCP1-KO mice, IL6 is strongly increased together with many other transcripts of inflammation associated genes (Kazak et al., 2017). Surprisingly, even the presence of UCP1 in preadipocytes alone was sufficient to bring about similar changes in the adipocytes as FGF2 pretreatment or expression. At least part of FGF2-programmed changes in preadipocytes finally leading to altered inflammatory state in resulting adipocytes is thus mediated through UCP1 in a necessary and sufficient manner.

The function of UCP1 in preadipocytes has been enigmatic since its first discovery. In the absence of adrenergic receptors and of significant lipid stores to mobilize, a classical thermogenic function through activation by free fatty acids appears unlikely. On the other hand, UCP1 is *per se* functional in this setting and uncouples respiration when activated. To be an active component in any mechanism, we must assume activators different from free fatty acids and an uncoupling function not primarily serving thermogenesis. In one such scenario, proposed earlier and supported by multiple reports, UCP1 may serve to prevent excessive inner membrane potential and/or respiratory chain redox pressure and thereby defend against reactive oxygen species (Echtay et al., 2003; Jastroch, 2017; Mailloux & Harper, 2011; Oelkrug et al., 2014). The latter and their products have been reported to activate UCP1 (Echtay et al., 2002, 2003; Malingriaux et al., 2013). Similarly, redox dependent UCP1 protein modification has been observed to regulate its activity (Chouchani et al., 2016) and, intriguingly, also inflammatory processes (Mills et al., 2022). Plausibly, such non-thermogenic functions of UCP1 are less or non-important in brown or beige adipocytes, but a key feature in UCP1-positive preadipocytes, a conjecture that remains to be tested.

Collectively, our findings demonstrate paracrine FGF2 signaling between preadipocytes to induce non-canonical UCP1 protein expression. By an unresolved UCP1-dependent mechanism, transcriptional regulation of inflammation-related genes is negatively programmed to finally give rise to mature adipocytes with decreased inflammatory status, IL6 and beta interferon associated gene products. UCP1-positive preadipocytes thus constitute a novel model to study endogenous, non-thermogenic UCP1 protein function. Elucidating this process and its molecular underpinnings will provide potential targets for the manipulation of adipose tissue inflammatory status.

## Acknowledgements and Funding Information

This research was funded by the German Research Foundation (Deutsche Forschungsgemeinschaft, DFG, Transregional Collaborative Research Center “BATenergy”: TRR333/1, project 450149205).

## Data availability

All underlying data are available upon request.

---

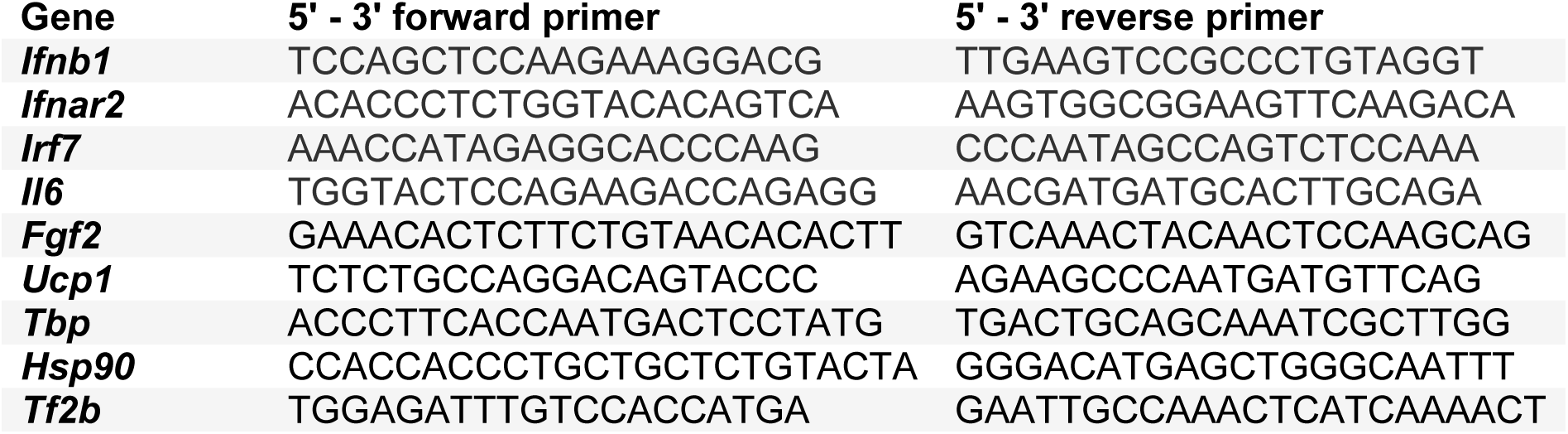

## Literature

Chan, C. C., Damen, M. S. M. A., Moreno-Fernandez, M. E., Stankiewicz, T. E., Cappelletti, M., Alarcon, P. C., Oates, J. R., Doll, J. R., Mukherjee, R., Chen, X., Karns, R., Weirauch, M. T., Helmrath, M. A., Inge, T. H., & Divanovic, S. (2020). Type I interferon sensing unlocks dormant adipocyte inflammatory potential. Nature Communications, 11(1), 2745. 10.1038/s41467-020-16571-4

Chen, A., Liao, S., Cheng, M., Ma, K., Wu, L., Lai, Y., Qiu, X., Yang, J., Xu, J., Hao, S., Wang, X., Lu, H., Chen, X., Liu, X., Huang, X., Li, Z., Hong, Y., Jiang, Y., Peng, J., … Wang, J. (2022). Spatiotemporal transcriptomic atlas of mouse organogenesis using DNA nanoball-patterned arrays. Cell, 185(10), 1777–1792.e21. 10.1016/j.cell.2022.04.003

Chouchani, E. T., Kazak, L., Jedrychowski, M. P., Lu, G. Z., Erickson, B. K., Szpyt, J., Pierce, K. A., Laznik-Bogoslavski, D., Vetrivelan, R., Clish, C. B., Robinson, A. J., Gygi, S. P., & Spiegelman, B. M. (2016). Mitochondrial ROS regulate thermogenic energy expenditure and sulfenylation of UCP1. Nature, 532(7597), Article 7597. 10.1038/nature17399

Dawid, C., Weber, D., Musiol, E., Janas, V., Baur, S., Lang, R., & Fromme, T. (2020). Comparative assessment of purified saponins as permeabilization agents during respirometry. Biochimica et Biophysica Acta (BBA) - Bioenergetics, 1861(10), 148251. 10.1016/j.bbabio.2020.148251

Echtay, K. S., Esteves, T. C., Pakay, J. L., Jekabsons, M. B., Lambert, A. J., Portero-Otín, M., Pamplona, R., Vidal-Puig, A. J., Wang, S., Roebuck, S. J., & Brand, M. D. (2003). A signalling role for 4-hydroxy-2-nonenal in regulation of mitochondrial uncoupling. The EMBO Journal, 22(16), 4103–4110. 10.1093/emboj/cdg412

Echtay, K. S., Roussel, D., St-Pierre, J., Jekabsons, M. B., Cadenas, S., Stuart, J. A., Harper, J. A., Roebuck, S. J., Morrison, A., Pickering, S., Clapham, J. C., & Brand, M. D. (2002). Superoxide activates mitochondrial uncoupling proteins. Nature, 415(6867), 96–99. 10.1038/415096a

Gantert, T., Henkel, F., Wurmser, C., Oeckl, J., Fischer, L., Haid, M., Adamski, J., Esser-von Bieren, J., Klingenspor, M., & Fromme, T. (2021). Fibroblast growth factor induced *Ucp1* expression in preadipocytes requires PGE2 biosynthesis and glycolytic flux. The FASEB Journal, 35(5). 10.1096/fj.202002795R

Gerngross, C., Schretter, J., Klingenspor, M., Schwaiger, M., & Fromme, T. (2017). Active brown fat during 18FDG-PET/CT imaging defines a patient group with characteristic traits and an increased probability of brown fat redetection. J Nucl Med. 10.2967/jnumed.116.183988

Han, M. S., White, A., Perry, R. J., Camporez, J.-P., Hidalgo, J., Shulman, G. I., & Davis, R. J. (2020). Regulation of adipose tissue inflammation by interleukin 6. Proceedings of the National Academy of Sciences, 117(6), 2751–2760. 10.1073/pnas.1920004117

Hu, C., Li, T., Xu, Y., Zhang, X., Li, F., Bai, J., Chen, J., Jiang, W., Yang, K., Ou, Q., Li, X., Wang, P., & Zhang, Y. (2023). CellMarker 2.0: An updated database of manually curated cell markers in human/mouse and web tools based on scRNA-seq data. Nucleic Acids Research, 51(D1), D870–D876. 10.1093/nar/gkac947

Jastroch, M. (2017). Uncoupling protein 1 controls reactive oxygen species in brown adipose tissue. Proceedings of the National Academy of Sciences, 114(30), 7744–7746. 10.1073/pnas.1709064114

Kazak, L., Chouchani, E. T., Stavrovskaya, I. G., Lu, G. Z., Jedrychowski, M. P., Egan, D. F., Kumari, M., Kong, X., Erickson, B. K., Szpyt, J., Rosen, E. D., Murphy, M. P., Kristal, B. S., Gygi, S. P., & Spiegelman, B. M. (2017). UCP1 deficiency causes brown fat respiratory chain depletion and sensitizes mitochondria to calcium overload-induced dysfunction. Proceedings of the National Academy of Sciences, 114(30), 7981–7986. 10.1073/pnas.1705406114

Kissig, M., Ishibashi, J., Harms, M. J., Lim, H., Stine, R. R., Won, K., & Seale, P. (2017). PRDM16 represses the type I interferon response in adipocytes to promote mitochondrial and thermogenic programing. The EMBO Journal, 36(11), 1528–1542. 10.15252/embj.201695588

Koledova, Z., Sumbal, J., Rabata, A., De La Bourdonnaye, G., Chaloupkova, R., Hrdlickova, B., Damborsky, J., & Stepankova, V. (2019). Fibroblast Growth Factor 2 Protein Stability Provides Decreased Dependence on Heparin for Induction of FGFR Signaling and Alters ERK Signaling Dynamics. Frontiers in Cell and Developmental Biology, 7, 331. 10.3389/fcell.2019.00331

Kowalczyk, M. S., Tirosh, I., Heckl, D., Rao, T. N., Dixit, A., Haas, B. J., Schneider, R. K., Wagers, A. J., Ebert, B. L., & Regev, A. (2015). Single-cell RNA-seq reveals changes in cell cycle and differentiation programs upon aging of hematopoietic stem cells. Genome Research, 25(12), 1860–1872. 10.1101/gr.192237.115

Kurosu, H., Ogawa, Y., Miyoshi, M., Yamamoto, M., Nandi, A., Rosenblatt, K. P., Baum, M. G., Schiavi, S., Hu, M.-C., Moe, O. W., & Kuro-o, M. (2006). Regulation of Fibroblast Growth Factor-23 Signaling by Klotho. Journal of Biological Chemistry, 281(10), 6120–6123. 10.1074/jbc.C500457200

Lundh, M., Pluciñska, K., Isidor, M. S., Petersen, P. S. S., & Emanuelli, B. (2017). Bidirectional manipulation of gene expression in adipocytes using CRISPRa and siRNA. Molecular Metabolism, 6(10), 1313–1320. 10.1016/j.molmet.2017.07.001

Mailloux, R. J., & Harper, M.-E. (2011). Uncoupling proteins and the control of mitochondrial reactive oxygen species production. Free Radical Biology and Medicine, 51(6), Article 6. 10.1016/j.freeradbiomed.2011.06.022

Malingriaux, E. A., Rupprecht, A., Gille, L., Jovanovic, O., Jezek, P., Jaburek, M., & Pohl, E. E. (2013). Fatty Acids are Key in 4-Hydroxy-2-Nonenal-Mediated Activation of Uncoupling Proteins 1 and 2. PLoS ONE, 8(10), Article 10. 10.1371/journal.pone.0077786

Matthias, A., Ohlson, K. B. E., Fredriksson, J. M., Jacobsson, A., Nedergaard, J., & Cannon, B. (2000). Thermogenic Responses in Brown Fat Cells Are Fully UCP1-dependent. Journal of Biological Chemistry, 275(33), 25073–25081. 10.1074/jbc.M000547200

Mills, E. L., Harmon, C., Jedrychowski, M. P., Xiao, H., Gruszczyk, A. V., Bradshaw, G. A., Tran, N., Garrity, R., Laznik-Bogoslavski, D., Szpyt, J., Prendeville, H., Lynch, L., Murphy, M. P., Gygi, S. P., Spiegelman, B. M., & Chouchani, E. T. (2022). Cysteine 253 of UCP1 regulates energy expenditure and sex-dependent adipose tissue inflammation. Cell Metabolism, 34(1), 140–157.e8. 10.1016/j.cmet.2021.11.003

Oeckl, J., Bast-Habersbrunner, A., Fromme, T., Klingenspor, M., & Li, Y. (2020). Isolation, Culture, and Functional Analysis of Murine Thermogenic Adipocytes. STAR Protocols, 1(3), 100118. 10.1016/j.xpro.2020.100118

Oelkrug, R., Goetze, N., Meyer, C. W., & Jastroch, M. (2014). Antioxidant properties of UCP1 are evolutionarily conserved in mammals and buffer mitochondrial reactive oxygen species. Free Radical Biology and Medicine, 77, 210–216. 10.1016/j.freeradbiomed.2014.09.004

Ornitz, D. M., & Itoh, N. (2015). The Fibroblast Growth Factor signaling pathway. WIREs Developmental Biology, 4(3), 215–266. 10.1002/wdev.176

Ornitz, D. M., & Itoh, N. (2022). New developments in the biology of fibroblast growth factors. WIREs Mechanisms of Disease, 14(4), e1549. 10.1002/wsbm.1549

Ornitz, D. M., Xu, J., Colvin, J. S., McEwen, D. G., MacArthur, C. A., Coulier, F., Gao, G., & Goldfarb, M. (1996). Receptor Specificity of the Fibroblast Growth Factor Family. Journal of Biological Chemistry, 271(25), 15292–15297. 10.1074/jbc.271.25.15292

Plotnikov, A. N., Hubbard, S. R., Schlessinger, J., & Mohammadi, M. (2000). Crystal Structures of Two FGF-FGFR Complexes Reveal the Determinants of Ligand-Receptor Specificity. Cell, 101(4), 413–424. 10.1016/S0092-8674(00)80851-X

Radványi, Á., & Röszer, T. (2024). Interleukin-6: An Under-Appreciated Inducer of Thermogenic Adipocyte Differentiation. International Journal of Molecular Sciences, 25(5), Article 5. 10.3390/ijms25052810

Rial, E. (1999). Retinoids activate proton transport by the uncoupling proteins UCP1 and UCP2. The EMBO Journal, 18(21), 5827–5833. 10.1093/emboj/18.21.5827

Ricquier, D. (2011). Uncoupling Protein 1 of Brown Adipocytes, the Only Uncoupler: A Historical Perspective. Frontiers in Endocrinology, 2. 10.3389/fendo.2011.00085

Seale, P., Bjork, B., Yang, W., Kajimura, S., Chin, S., Kuang, S., Scimè, A., Devarakonda, S., Conroe, H. M., Erdjument-Bromage, H., Tempst, P., Rudnicki, M. A., Beier, D. R., & Spiegelman, B. M. (2008). PRDM16 controls a brown fat/skeletal muscle switch. Nature, 454(7207), 961–967. 10.1038/nature07182

Shamsi, F., Xue, R., Huang, T. L., Lundh, M., Liu, Y., Leiria, L. O., Lynes, M. D., Kempf, E., Wang, C.-H., Sugimoto, S., Nigro, P., Landgraf, K., Schulz, T., Li, Y., Emanuelli, B., Kothakota, S., Williams, L. T., Jessen, N., Pedersen, S. B., … Tseng, Y.-H. (2020). FGF6 and FGF9 regulate UCP1 expression independent of brown adipogenesis. Nature Communications, 11(1), 1421. 10.1038/s41467-020-15055-9

Urakawa, I., Yamazaki, Y., Shimada, T., Iijima, K., Hasegawa, H., Okawa, K., Fujita, T., Fukumoto, S., & Yamashita, T. (2006). Klotho converts canonical FGF receptor into a specific receptor for FGF23. Nature, 444(7120), 770–774. 10.1038/nature05315

Virtanen, K. A., Lidell, M. E., Orava, J., Heglind, M., Westergren, R., Niemi, T., Taittonen, M., Laine, J., Savisto, N. J., Enerback, S., & Nuutila, P. (2009). Functional brown adipose tissue in healthy adults. N Engl J Med, 360(15), Article 15. 10.1056/NEJMoa0808949

Wang, H., Willershäuser, M., Karlas, A., Gorpas, D., Reber, J., Ntziachristos, V., Maurer, S., Fromme, T., Li, Y., & Klingenspor, M. (2019). A dual Ucp1 reporter mouse model for imaging and quantitation of brown and brite fat recruitment. Molecular Metabolism, 20, 14–27. 10.1016/j.molmet.2018.11.009

Westphal, S., Gantert, T., Kless, C., Hüttinger, K., Klingenspor, M., & Fromme, T. (2019). Fibroblast growth factor 8b induces uncoupling protein 1 expression in epididymal white preadipocytes. Scientific Reports, 9(1), 8470. 10.1038/s41598-019-44878-w

Wu, J., Boström, P., Sparks, L. M., Ye, L., Choi, J. H., Giang, A.-H., Khandekar, M., Virtanen, K. A., Nuutila, P., Schaart, G., Huang, K., Tu, H., van Marken Lichtenbelt, W. D., Hoeks, J., Enerbäck, S., Schrauwen, P., & Spiegelman, B. M. (2012). Beige Adipocytes Are a Distinct Type of Thermogenic Fat Cell in Mouse and Human. Cell, 150(2), 366–376. 10.1016/j.cell.2012.05.016

Yayon, A., Klagsbrun, M., Esko, J. D., Leder, P., & Ornitz, D. M. (1991). Cell surface, heparin-like molecules are required for binding of basic fibroblast growth factor to its high affinity receptor. Cell, 64(4), 841–848. 10.1016/0092-8674(91)90512-W

Yeh, B. K., Igarashi, M., Eliseenkova, A. V., Plotnikov, A. N., Sher, I., Ron, D., Aaronson, S. A., & Mohammadi, M. (2003). Structural basis by which alternative splicing confers specificity in fibroblast growth factor receptors. Proceedings of the National Academy of Sciences, 100(5), 2266–2271. 10.1073/pnas.0436500100

